# Enhanced loss of retinoic acid network genes in *Xenopus laevis* achieves a tighter signal regulation

**DOI:** 10.1101/2022.01.04.474867

**Authors:** Tali Abbou, Liat Bendelac-Kapon, Audeliah Sebag, Abraham Fainsod

## Abstract

Retinoic acid (RA) is a major regulatory signal during embryogenesis produced from vitamin A (retinol) by an extensive, autoregulating metabolic and signaling network to prevent fluctuations that result in developmental malformations. *Xenopus laevis* is an allotetraploid hybrid frog species whose genome includes L (long) and S (short) chromosomes from the originating species. Evolutionarily, the *X. laevis* subgenomes have been losing either L or S homoeologs in about 43% of genes to generate singletons. In the RA network, out of the 47 genes, about 46% have lost one of the homoeologs, like the genome average. In contrast, RA metabolism genes from storage (retinyl esters) to retinaldehyde production exhibit enhanced gene loss with 75% singletons out of 28 genes. The effect of this gene loss on RA signaling autoregulation was studied. Employing transient RA manipulations, homoeolog gene pairs were identified in which one homeolog exhibits enhanced responses or looser regulation than the other, while in other pairs both homoeologs exhibit similar RA responses. CRISPR /Cas9 targeting of individual homoeologs to reduce their activity supports the hypothesis where the RA metabolic network gene loss results in tighter network regulation and more efficient RA robustness responses to overcome complex regulation conditions.

## 1. Introduction

Retinoic acid (RA) signaling is one of the major regulatory pathways active during embryogenesis and it controls numerous developmental processes [1–4]. RA is produced in the body from nutritional sources containing vitamin A (retinol, ROL) and other retinoids, or carotenoids [5,6]. This metabolic network involves multiple enzymes active in the conversion of these substrates into RA, or the storage of retinoids as retinyl esters to survive periods when the nutritional sources of retinoids or carotenoids are diminished or lacking [5,7–10]. Increased or decreased RA signaling levels can result in severe developmental malformations [8,11–14]. For this reason, the RA metabolic and signaling network exhibits efficient self-regulation to overcome fluctuations in this signaling pathway elicited by dietary changes, environmental toxicity, or genetic polymorphisms [15–26]. The autoregulation of the RA network by RA levels is instrumental to maintaining pathway robustness in response to hampering mechanisms [27,28].

For over half a century, gene duplication has been studied as one of the processes important for the evolution of species [29–31]. Following duplication of a gene or gene family, one of the copies can be lost, or one member retains the original function and regulation whereas the extra copy can gain novel functions, called neofunctionalization and regulation. Generally, duplications occur on a small scale involving restricted genomic regions, but in extreme cases, gene duplication of all the genes present in the genome is the result of polyploidization by whole-genome duplication (WGD) [29]. *Xenopus laevis*, the African clawed frog, was proposed to have arisen by interspecific hybridization of two related diploid species followed by WGD making it an allotetraploid species [32]. From sequence analysis of the *Xenopus laevis* genome, it was concluded that the two ancestral originating species separated about 34 million years ago and the allotetraploid event occurred about 17-18 million years ago. *X. laevis* inherited half of its genome from each ancestor, resulting in two distinct subgenomes present in a diploid condition: L and S, for long and short chromosomes, respectively [32].

As a result of a WGD event, one of the gene copies may diverge, probably together with other genes, co-evolving as a complex system to achieve neofunctionalization or subfunctionalization [31]. Alternatively, it has been proposed that sometimes following a burst of increase in genome complexity, there is a long process of genome reduction [33,34]. During the genome reduction process following a WGD event, one of the gene copies might be lost or mutated into a pseudogene [35]. Analysis of the genome of *X. laevis* and its transcriptome clearly identified many instances of the L and S genes and their respective transcripts [32,36–39]. Also important, these studies identified a process of gene loss that encompasses today about 43% of protein-coding genes [32]. Most of this gene loss took place in the S subgenome. Detailed genome analysis also revealed high (>85%) conservation of both (L and S) gene copies, i.e., homoeologs, among genes encoding DNA binding proteins, transcription factors, the Wnt, Hh, Notch, FGF, TGFß, and Hippo signaling pathways [32,40–42]. The inverse situation, enhanced gene loss, was observed among the genes encoding DNA repair proteins (79% singletons). This high rate of singletons was attributed to either a lack of selective pressure where one enzyme encoding locus is sufficient or to functional incompatibility among homoeologs leading to deletions [32].

We analyzed the RA signaling network in *X. laevis* and uncovered an average distribution of homoeologs and singletons compared to the genomic average. However, deeper analysis of the RA metabolic and gene regulatory network revealed that among the enzymatic components necessary for retinaldehyde (RAL) production, and retinoid storage and retrieval there is a high incidence of gene loss, resulting in about 75% singletons. For the rest of the network, from RAL to RA production, RA disposal and gene regulation, most genes (about 95%), are present as homoeolog pairs. We studied the possibility that the enhanced gene loss in the RA metabolic network leading to RAL production is related to the regulation of the RA signal to prevent teratogenic outcomes. A transient RA manipulation approach followed by kinetic analysis of the recovery period revealed the presence of both tightly regulated (restricted responses, low RA fluctuation sensitivity) and loosely regulated (enhanced responses, high RA change sensitivity) genes. We speculated that homoeolog pairs with markedly different RA responses would degrade the robustness response to RA fluctuations, therefore we used CRISPR / Cas9 to target specific homoeologs. Our results showed that among homoeolog pairs with similar RA responses, individual knockdowns resulted in similar recovery kinetics from the RA treatment. In contrast among homoeologs with diverged RA responses, knockdown of the tightly-regulated homoeolog impairs the kinetic recovery response, whereas, targeting the loosely regulated homoeolog improves the RA robustness response. These results support our hypothesis proposing that enhanced gene loss of the RA network components might lead to an improved robustness response by reducing the number of genes to be regulated, specifically removing genes with enhanced responses to RA fluctuation.

## 2. Materials and Methods

### 2.1. Embryo Culture and Treatment

*Xenopus laevis* frogs were purchased from Xenopus I or NASCO (Dexter, MI or Fort Atkinson, WI, United States). Experiments were performed after approval and under the supervision of the Institutional Animal Care and Use Committee (IACUC) of the Hebrew University (Ethics approval no. MD-17-15281-3). Embryos were obtained by *in vitro* fertilization, incubated in 0.1% Modified Barth’s Solution and Hepes (MBSH), and staged according to Nieuwkoop and Faber [43]. All-*trans* retinoic acid and Dimethyl sulfoxide (DMSO) were purchased from Sigma-Aldrich (St. Louis, MO, United States). Stock solutions of RA were prepared in DMSO. For transient RA treatment, embryos were placed in 10 nM or 25 nM RA from late blastula (st. 9.5) and washed two hours later, at early gastrula (st. 10.25) by three changes of 0.1% MBSH and further incubated in fresh 0.1% MBSH for the desired time. Samples were collected 1, 1.5, 2, and 2.5 hours after washing.

### 2.2. Quantitative reverse transcription real-time PCR (qPCR)

Total RNA from embryos was extracted with Aurum™ Total RNA Mini Kit (Bio-Rad). cDNA was synthesized using iScript cDNA Synthesis Kit (Bio-Rad). The real-time PCR reactions were performed using the CFX384 Real Time System (Bio-Rad) and iTaq Universal SYBR Green Supermix (Bio-Rad). Each experiment was repeated at least three independent times and each time the samples were run in triplicate. *slc35b1*.L was used as the housekeeping reference gene. The primers used for qPCR analysis are listed in Table 1.

**Table 1.**
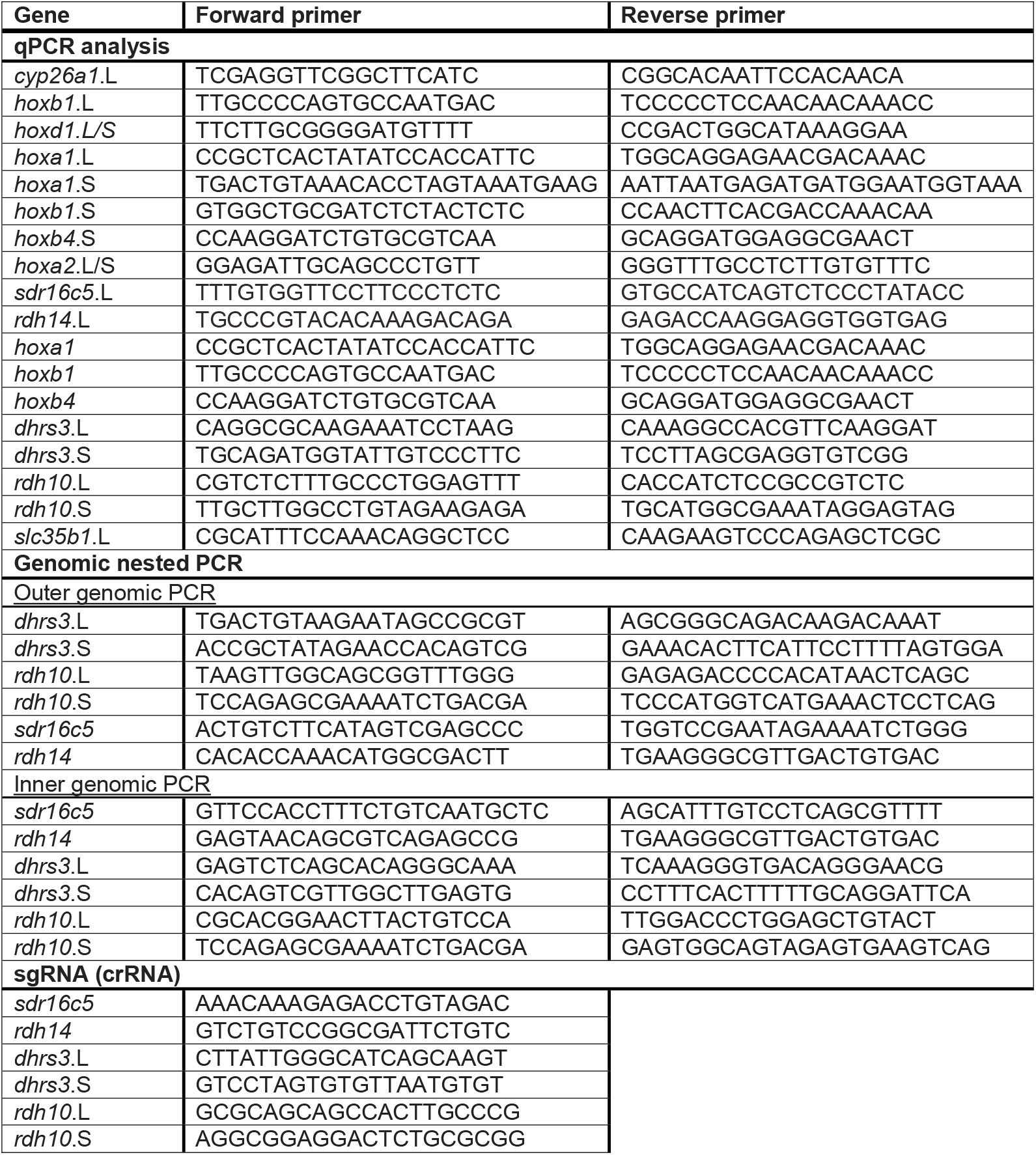
Sequences for qPCR and genomic DNA amplification primers and sgRNAs.

### 2.3. Generation of CRISPant embryos

For gene-specific single guide RNA design (sgRNA), genomic DNA sequences were selected from Xenbase.org [44] for the L and S homoeologs when present and analyzed using CRISPRdirect [45] and CRISPRscan [46] for target site search. Computational estimation of the sgRNA efficiency was determined using the inDelphi software [47,48]. For the generation of F0 CRISPant embryos, we injected one-cell stage embryos with Cas9 ribonucleoprotein (RNP) complexes employing the two-RNA component (crRNA:tracrRNA) approach [49]. Briefly, chemically synthesized and modified for stability (Alt-R) RNAs (crRNA and tracrRNA; IDT, USA) (Table 1) were annealed to generate the double guide complexes (crRNA:tracrRNA) and were incubated (10 min at 37°C) with *S. pyogenes* Cas9 protein (IDT, USA) to generate RNP complexes. Eight nanoliters of the RNP complex solution were injected into the cytoplasm of one-cell stage embryos.

To determine the efficiency of indel induction, genomic DNA was extracted from 5 individual embryos at mid-gastrula (st. 11) or later employing the GenElute Mammalian Genomic DNA Miniprep Kit (Sigma). The genomic region containing the CRISPR / Cas9 targeted region was PCR amplified using a nested PCR approach (Table 1) and the size-selected and cleaned product was sequenced. Genome editing efficiency was analyzed by decomposition analysis [50] using the Synthego ICE algorithm [51].

### 2.4. Statistical analysis

All statistical comparisons were carried out using the Prism software package (Graph Pad Software Inc., San Diego, CA). Results are given as the mean ± standard error of the mean (SEM). Tests used were the 2-tailed t-test for two-sample comparisons, Dunnett’s (ANOVA) multiple comparisons test, or Fisher test. Differences between means were considered significant at a significance level of p<0.05.

## 3. Results and Discussion

### 3.1. Conservation of the RA network

In a recent extensive data mining effort searching the KEGG and Xenbase databases [44,52] and the literature, we assembled the putative components of the RA metabolic and signaling network during early embryogenesis [1,6,9]. To determine the composition of the RA network during gastrula in *Xenopus laevis* embryos we assessed which components are expressed during this developmental stage based on analysis of transcriptomic data sets [27,53,54] and corroborated the gastrula expression by quantitative RT-PCR (qPCR) [55]. This view of the RA metabolic and signaling network exhibits a rather uncommon characteristic that for each enzymatic step, multiple genes encoding enzymes have been described that are capable of performing the same reaction (Fig. 1). For some of the reversible reactions, such as the oxidation of ROL to RAL and the corresponding reduction of RAL to ROL, a preferred activity has been identified, but the reverse reaction might be possible under certain conditions [56–58]. Because it is necessary to maintain non-teratogenic levels of RA at these stages, there appear to be several ways to control levels of RA signaling. Substrate availability for the retinaldehyde dehydrogenase enzymes that oxidize RAL to RA, the ALDH1A subfamily, is controlled by the presence of multiple enzymes that reduce RAL back to ROL [10,13,15,25,59–61]. Levels of signaling can additionally be controlled by regulating the spatial-temporal expression of the many RA metabolic and gene regulatory network components, including the RA nuclear receptor families (RAR and RXR) and retinoid-binding proteins [62–64].

**Figure 1.**
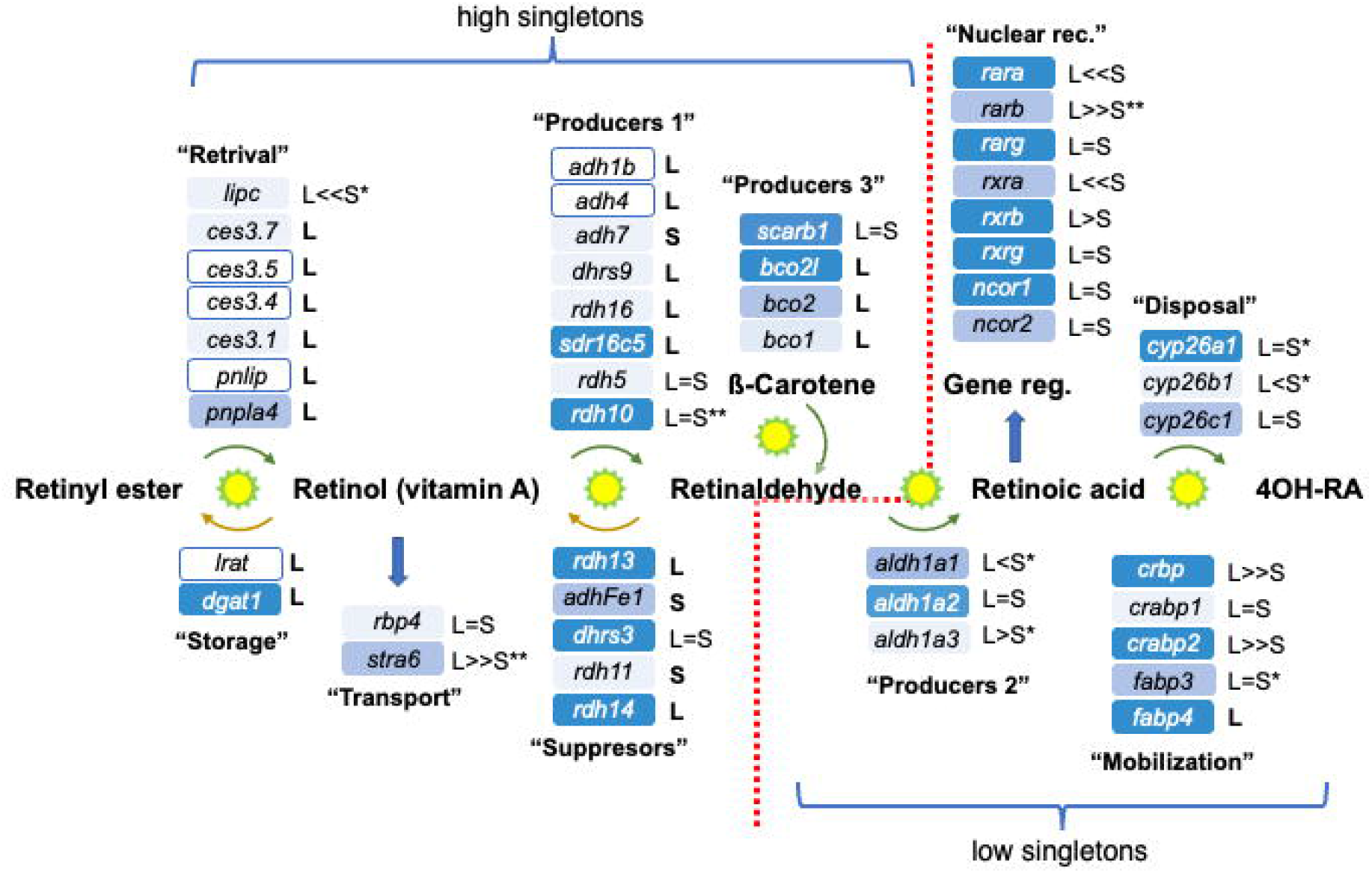
Evolutionary conservation of the RA metabolic and gene-regulatory network genes in *Xenopus laevis*. Composition of the RA metabolic and gene-regulatory network in the *Xenopus laevis* genome based on KEGG and Xenbase database analysis and literature searches [44,52]. Expression of the RA network components during gastrula stages was determined from our own and published transcriptomic datasets. The relative expression shown (blue shades) during gastrula stages is based on Session et al., [32]. Dark blue, •10 TPM; mid blue, >0.5-<10 TPM; light blue, •0.5 TPM; white, no data in the transcriptomic dataset. The homoeolog /singleton status of each gene is marked (L and /or S). The relative expression levels between homoeologs are summarized: =, similar expression levels; <, 3-6 fold difference; <<, more than 6 fold difference. Asterisks indicate whether temporal expression patterns of the homeologs are similar (no asterisk), partially divergent (*), or highly divergent (**).

Analysis of the allotetraploid status of the genes encoding all the identified RA network components in the *Xenopus laevis* genome revealed that in 25 out of 47 genes both the L and S homoeologs [32] have been retained during evolution (Fig. 1; Table 2). Thus, 22 genes (46.8%) in the RA network have lost one of the homoeologs, bringing this metabolic and signaling network close to the genomic average (43.6%) of singletons among protein-coding genes [32]. This observation seemingly contradicts previous studies of other main signaling pathways critical for normal embryogenesis, like TGFß, FGF, Wnt, Hh, Notch, Hippo, and of transcription factors for which the conservation of the L and S homoeologs is very high (>83.3%; Table 2)[40–42].

**Table 2.** Singleton distribution in *Xenopus laevis* metabolic and signaling pathways.

| Pathway | Total genes | Singletons | % Singletons |
| --- | --- | --- | --- |
| **High singleton** |  |  |  |
| Vitamin D from 7-dehydrocholesterol<sup>1</sup> | 9 | 6 | 66.7% |
| RA up to RAL<sup>1</sup> | 28 | 21 | 75% |
| Folic acid metabolism<sup>1</sup> | 27 | 19 | 70.4% |
| DNA repair | 57 | 45 | 78.9% |
| **Average genome singletons** |  |  |  |
| De novo Purine biosynthesis<sup>1</sup> | 15 | 7 | 46.7% |
| Thyroid hormone synthesis<sup>1</sup> | 52 | 23 | 44.2% |
| Protein-coding<sup>2</sup> | >13781 |  | 43.6% |
| RA signaling metabolism and signaling | 47 | 22 | 46.8% |
| Vitamin D incl. 7-dehydrocholesterol<sup>1</sup> | 21 | 8 | 38.1% |
| **Suppressed singletons** |  |  |  |
| Glycolysis/Gluconeogenesis/Krebs cycle<sup>1</sup> | 44 | 9 | 20.5% |
| Notch<sup>3</sup> | 48 | 8 | 16.7% |
| MicroRNAs<sup>2</sup> | 180 | 24 | 13.3% |
| Transcription Factors<sup>4</sup> | 218 | 28 | 12.8% |
| Wnt<sup>3</sup> | 108 | 13 | 12% |
| Hippo<sup>3</sup> | 48 | 5 | 10.4% |
| BMP/TGFβ<sup>5</sup> | 126 | 13 | 10.3% |
| RA from RAL<sup>1</sup> | 19 | 1 | 5.2% |
| FGF<sup>5</sup> | 60 | 3 | 5% |
| cis-regulatory elements (non-coding)<sup>2</sup> | 550 | 9 | 1.6% |
| Hh<sup>3</sup> | 18 | 0 | 0% |
| HSPG<sup>3</sup> | 16 | 0 | 0% |
| TLE<sup>3</sup> | 4 | 0 | 0% |

<sup>1</sup> Data from KEGG, this work; <sup>2</sup>Session et al., 2016; <sup>3</sup>Michiue et al., 2017; <sup>4</sup>Watanabe et al., 2017; <sup>5</sup>Suzuki et al., 2017.

Further analysis of the distribution of homoeologs and singletons within the RA network genes, however, revealed a surprising, non-random distribution of gene loss events (Fig. 1). Among the enzymes involved in the metabolic steps leading up to the production of RAL, including retinoid storage, about 75% of the genes (21 out of 28) were encoded as singletons (Fig. 1; Table 2). Interestingly, for genes involved in the oxidation of RAL to RA, hydroxylation of RA, or actual RA-driven gene regulation, almost all (94.8%; 18 out of 19) are still encoded by both L and S homoeologs (Fig. 1; Table 2).

This suggests a preferential loss of homoeologs involved in the production of RAL, regulation of RAL production, or storage of retinoids. One possible explanation for this asymmetrical gene loss in the metabolic side of the RA network could be the observation that RAL availability for oxidation by retinaldehyde dehydrogenases is like a “commitment” step (Fig. 1). The oxidation of RAL to RA by the ALDH1A1, ALDH1A2, and ALDH1A3 enzymes cannot be reversed and either promotes RA-driven gene regulation or the RA produced has to be neutralized and degraded. RA signaling is one of the major embryonic signaling pathways dependent on the maternal nutritional status and its function can be altered by environmental factors [5,15–17]. Fluctuations in RA signaling, increase or decrease, can be extremely teratogenic. Therefore, we explored the possibility that the extensive gene loss observed preferentially achieves tighter regulation of the RA signal, providing an evolutionary advantage.

To assess whether this preferential gene loss in the RA metabolic and gene-regulatory pathway was restricted to RA signaling, we also analyzed the genomic evolution of two additional nuclear receptor signaling pathways closely linked to RA: vitamin D and thyroid hormone signaling [65,66]. Thyroid hormone biosynthesis and signaling in *Xenopus* includes 23 genes out of 52 that are already singletons (44.2%), bringing this pathway to the genomic average with no obvious distinctive distribution (Table 2). Interestingly, vitamin D biosynthesis and gene regulation exhibit a distribution resembling the RA network where part of the pathway is rich in singletons (Fig. 2). Also, in this case, the whole pathway (21 genes) exhibits 38% singletons (8 genes) close to the protein-coding average (Table 2) [32]. But, from the production of cholecalciferol (Vitamin D_3_) the pathway has 6 genes out of 9 that are already singletons (Fig. 2; Table 2). These 66.7% singletons in the vitamin D-specific part of the network show a preferential loss of genes involved in metabolism and gene regulation by this ligand.

**Figure 2.**
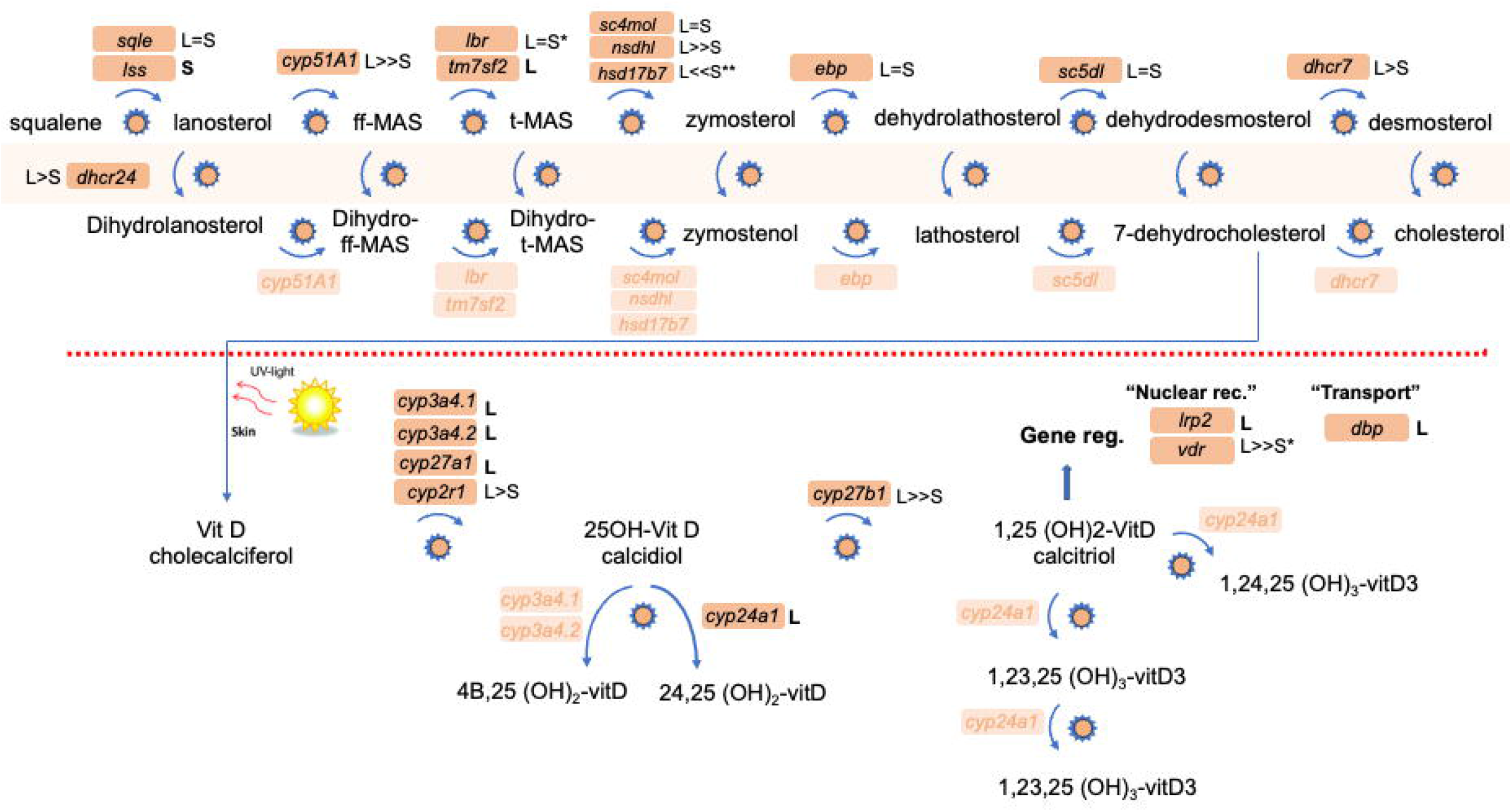
Homoeolog and singleton status of genes involved in Vitamin D metabolism and signaling. KEGG analysis of the vitamin D metabolic and signaling network identified 21 genes in the *X. laevis* genome. The homoeolog /singleton status of each gene is marked (L and /or S). In the metabolic part of the pathway leading to cholesterol production (above the red dotted line), the pathway runs in parallel from lenosterol and dehydrolanosterol. The relative expression levels between homoeologs are summarized: =, similar expression levels; <, 3-6 fold difference; <<, more than 6 fold difference. Asterisks indicate whether temporal expression patterns of the homeologs are similar (no asterisk), partially divergent (*), or highly divergent (**).

As both the RA and vitamin D signaling pathways involve biosynthesis of the regulatory ligand through a metabolic pathway, to assess the generality of this observation we explored the L or S gene loss in additional metabolic pathways. For the *de novo* purine biosynthesis pathway we scored 7 genes out of 15 that are already singletons (Table 2). The 47% singleton status is close to the genomic average suggesting the normal rate of gene loss for coding sequences. Analysis of glycolysis + gluconeogenesis + Krebs cycle identified 9 singletons among 44 genes (20.4%), which is a low singleton proportion suggesting conservation of both homoeologs in these metabolic pathways (Table 2). Analysis of the folic acid metabolic network indicated the reverse: a high proportion of singletons. From the information in KEGG, we identified 27 enzyme-encoding genes in the *X. laevis* genome out of which 19 (70.4%) are encoded by singletons (Table 2), indicating a preferential loss of one of the homoeologs during evolution.

Based on the analysis of homoeolog loss in the *Xenopus laevis* genome to date [32,40– 42], signaling pathways and metabolic networks can be classified into three groups. There are pathways that exhibit homoeolog retention rates similar to the protein-coding gene average in the whole genome. From our analysis, we identified the thyroid hormone synthesis and signaling, vitamin D biosynthesis and signaling, *de novo* purine biosynthesis, and RA metabolism and signaling pathways as belonging to this group (Table 2). The second group includes previously analyzed signaling pathways described as having very high homoeolog retention rates (low or suppressed singletons)[32,40– 42], and we added Krebs cycle, glycolysis, and gluconeogenesis to this group (Table 2). Our analysis identified the third group as having a high rate of gene loss creating a high proportion of singletons (Table 2). Interestingly, apart from the folate metabolic network, for both RA metabolism and vitamin D biosynthesis and signaling, the high rate of homoeolog loss localizes to a specific region of the pathway (Figs. 1 and 2).

### 3.2 Genomic changes in the loss of a homoeolog

The high incidence of singletons in the RA metabolic network from retinyl ester storage to the production of RAL prompted us to try to understand the genomic rearrangements that resulted in this enhanced homoeolog loss that involved 21 out of 28 genes (Table 2). We focused our analysis on enzymes with alcohol dehydrogenase activity to oxidize vitamin A to RAL (Producers 1); enzymes, mainly of the short-chain dehydrogenase / reductase family, that reduce RAL to retinol (Suppressors); and proteins involved in ß-carotene cleavage to RAL (Producers 3) (Fig. 1; Table 3). We compared the appropriate genomic regions between the L and S chromosomes choosing the first pair of conserved genes flanking the deleted or rearranged region as boundaries (Table 3). Using these flanking genes, we could determine the distance between them in the L and S chromosomes and analyze the region between them (Table 3). This type of analysis revealed cases in which single or multiple genes were deleted. Also, the length of the modified region changed from 0.1 to 410 Kb (Table 3). We could group the rearrangements into three groups. The first group represents singletons in which the loss of a homoeolog involved a relatively small (0.1-36 Kb) and simple deletion (Fig. 3A; Table 3). The second group involved large (102-410 Kb) deletions (Fig. 3B; Table 3). In the third group, we identified large deletions (81 and 238 Kb) combined with extensive rearrangement of the genomic region (Fig. 3C; Table 3). These results show that most deletions leading to the loss of a homoeolog are relatively simple although the genomic region lost can be small or very large. In a few cases, the loss of a homoeolog involved complex genomic rearrangements in addition to the deletion of genes. In the locus on chromosome 1 (Fig. 3C) there are multiple *adh* genes suggesting the possibility that this region contained duplicated sequences that could contribute to the rearrangements in this genomic region. On chromosome 9_10 we also observed a complex deletion but did not observe gene duplications that could contribute to its creation.

**Table 3.**
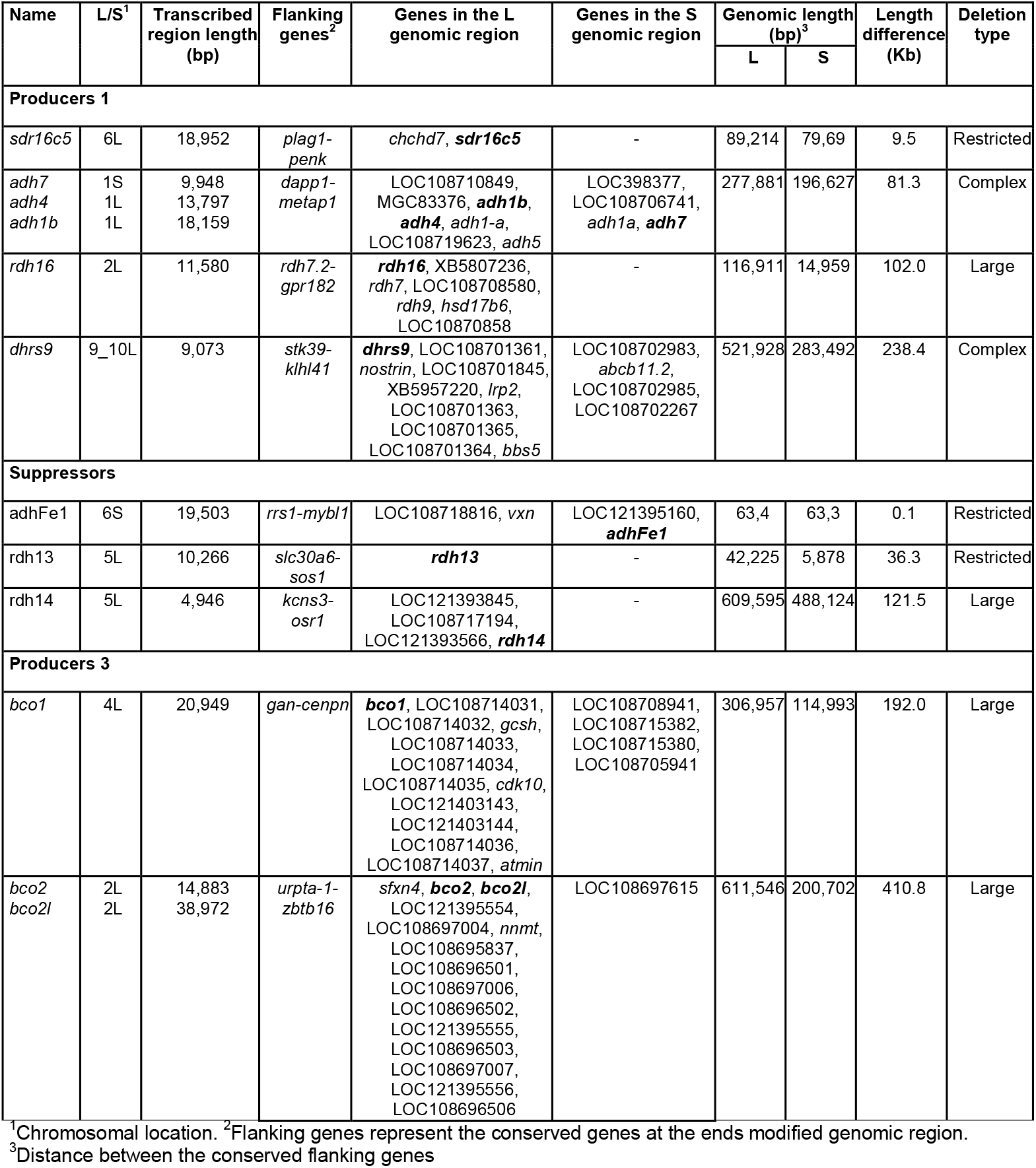
Genomic changes in the generation of RA network singletons.

**Figure 3.**
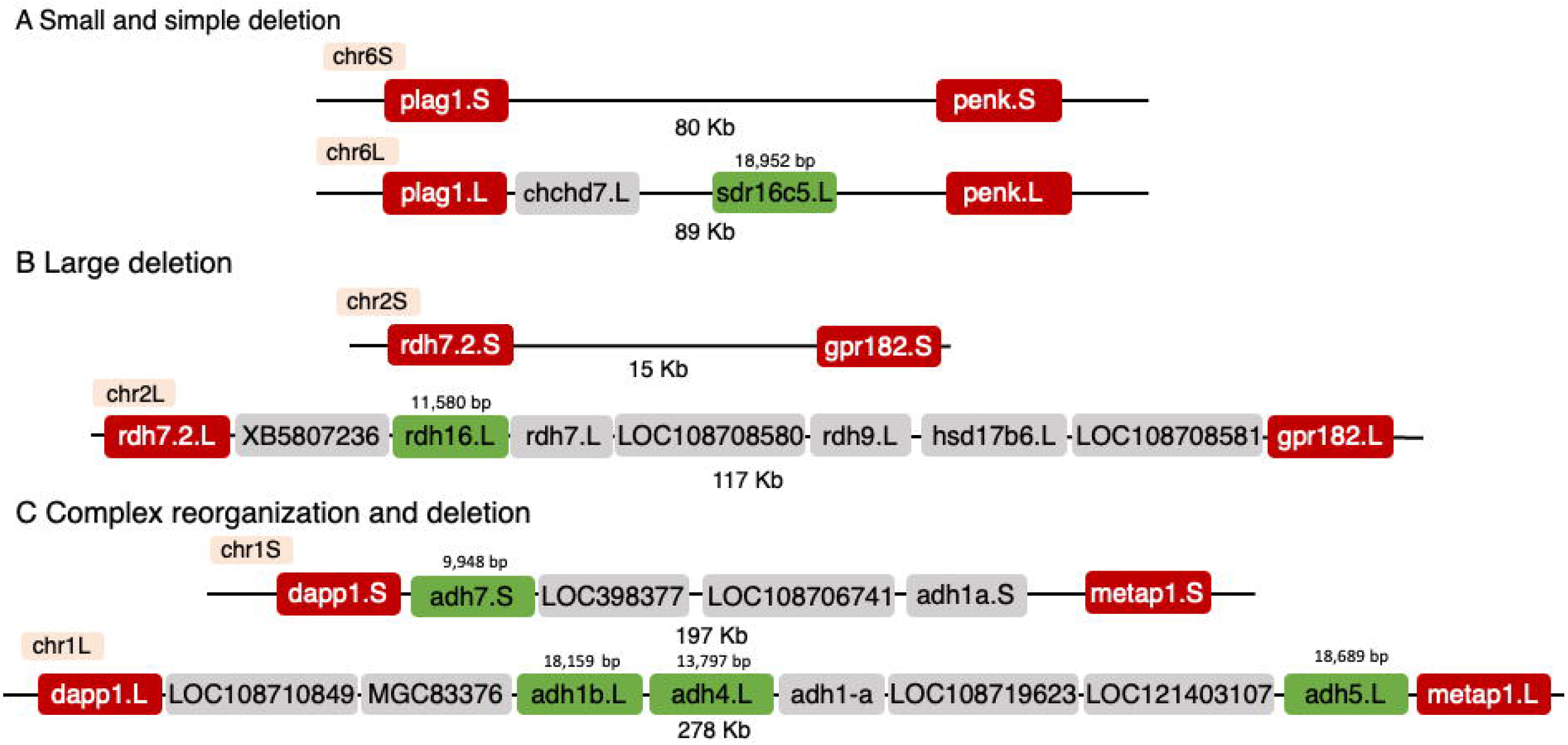
Genomic rearrangements involving RA network genes. Schematic examples of the types of genomic deletions and rearrangements observed in the deletion of homoeologs. The gene conserved, i.e., singleton, is marked in green. The flanking genes selected to determine the interval that was deleted are marked in red. Additional genes or putative coding sequences within the regions are marked in gray. (A) Generation of the *sdr16c5*.L singleton apparently involved the deletion of a small genomic region on chromosome 6S. (B) The deletion to create the *rdh16*.L singleton involved deleting about 100 Kb on chromosome 2S including multiple genes. (C) The genomic reorganization and deletion on chromosome 1 created the *adh7*.S singleton on chromosome 1S and singletons for *adh1*.L, *adh4*.L, and *adh5*.L on chromosome 1L.

### 3.3. Expression overlaps and responsiveness of RA network components

Our recent analysis of the RA metabolic and signaling network revealed a high degree of robustness following disruption of this pathway within the physiological range [27]. Moreover, this study showed that enzymes performing the same metabolic reaction and expressed in partially overlapping patterns might be regulated differently. These differential responses to RA fluctuation are part of the mechanism to keep this critical signal within an appropriate, non-teratogenic range [27]. One possibility for the preferential loss of homoeologs in genes encoding RA network components is the selective or non-selective reduction of gene copies to achieve tighter regulation of the signal. To begin exploring these possibilities as possible driving forces for gene loss, we searched for gene pairs expressed during gastrula stages that have the same enzymatic activity but one of them is a singleton and the other is still presented as a homoeolog pair. Based on our previous studies we chose *rdh10* and *sdr16c5* for genes encoding enzymes that oxidize ROL to RAL (Producers 1), *dhrs3* and *rdh14* for genes encoding enzymes that reduce RAL to ROL (Suppressors), and *aldh1a2* and *aldh1a3* for genes encoding enzymes that oxidize RAL to RA and for which no singletons are known (Producers 2) (Fig. 1). The temporal pattern of expression was determined for these genes including analysis of the individual homoeologs by qPCR (Fig. 4). In all three gene groups, we observed extensive overlap in the temporal expression pattern between the selected singleton and at least one of the homoeologs of the paired gene with similar enzymatic activity (Fig. 4A-C). While *sdr16c5* (singleton) exhibits a pattern similar to the *rdh10*.S homoeolog both having significant maternal expression (Fig. 4A), the *rdh10*.L homoeolog is mainly expressed zygotically. Among the Suppressors tested (Fig. 4B), *rdh14* exhibits what might be maternal transcripts, and its expression levels decline during gastrulation (Fig. 4B). Both *dhrs3* homoeologs retain extensively overlapping temporal expression patterns that peak at late gastrulation and subsequently decline (Fig. 4B). The retinaldehyde dehydrogenase encoding genes *aldh1a2* and *aldh1a3* exhibit mostly zygotic expression patterns with a marked up-regulation with the onset of gastrulation (Fig. 4C). These expression patterns support the partial overlap between the genes selected and are part of the RA network during gastrulation.

**Figure 4.**
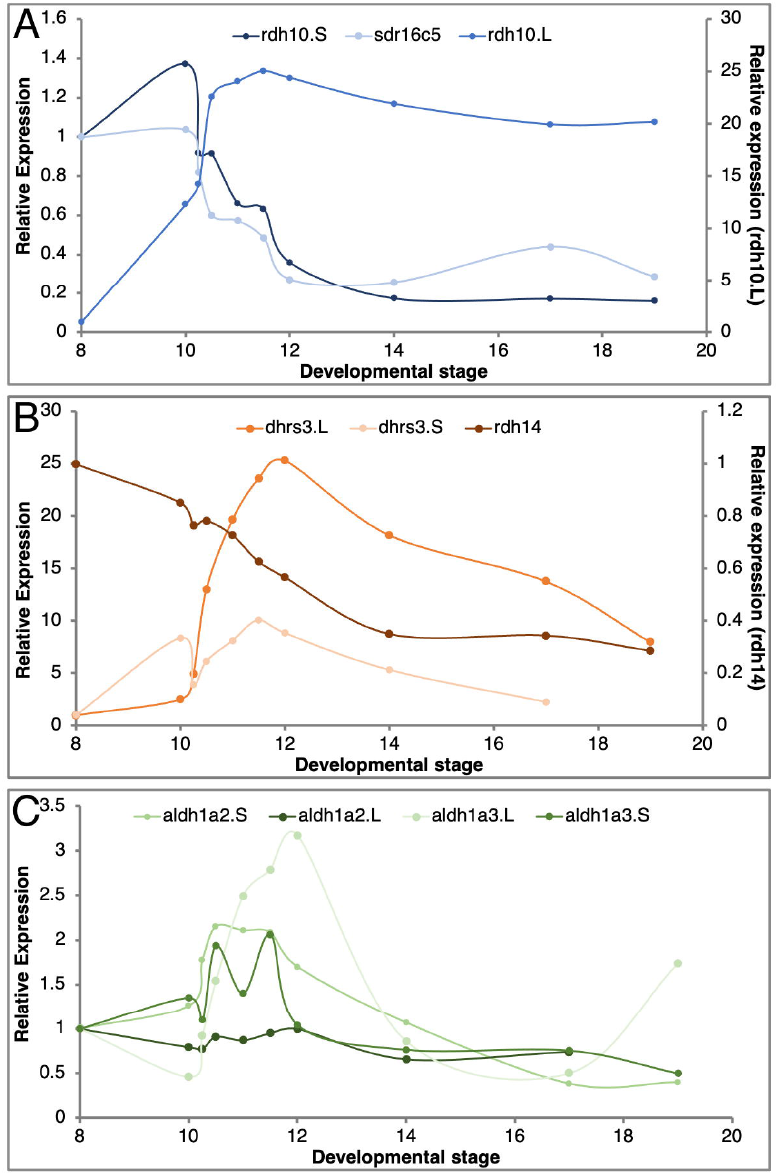
Comparative temporal expression pattern of homoeologs and singletons. Embryos were collected at different developmental stages from blastula to mid-neurula. The temporal expression pattern of each gene was determined by qPCR. (A) Expression of the *rdh10*.L, *rdh10*.S, and *sdr16c5*.L genes (Producers 1). (B) Temporal expression pattern of *dhrs3*.L, *dhrs3*.S and *rdh14*.L (Suppressors). (C) Expression pattern of *aldh1a2*.L, *aldh1a2*.S, *aldh1a3*.L and *aldh1a3*.S (Producers 2).

For the four homoeolog pairs studied, we observed divergence in their temporal expression patterns. In the extreme case, *rdh10*.S exhibits high levels of maternal transcripts and a gradual decline during gastrula stages, whereas *rdh10*.L expression is activated after the midblastula transition as a zygotic gene (Fig. 4A). A similar expression pattern for this homoeolog pair was determined from transcriptomic data [32]. This divergence in temporal expression patterns suggests changes in regulatory elements and initial subfunctionalization of the homoeologs [36]. For two of the other homoeolog pairs, *dhrs3* and *aldh1a3*, there are subtle differences in their temporal expression patterns, with extensive overlap but also new gene-specific changes (Fig. 4B,C). In contrast, the *aldh1a2* homoeologs exhibit temporal expression patterns that are very similar (Fig. 4C). Interestingly, the early expression of *aldh1a2* at the onset of gastrulation has been linked to the onset of RA signaling as the enzyme encoded by this gene is the last component needed to complete the biosynthesis of RA [15,67–70]. Also, we have shown that within the *aldh1a* gene family, *aldh1a2* is expressed at the highest levels during early gastrula stages [15]. Thus, there appears to be selective pressure to conserve this expression pattern and the early gastrula activity of *aldh1a2*.

To better understand the contribution of these genes to the response of the RA network to fluctuations in ligand levels, we studied the response of these genes to subtle manipulation of RA levels. Analysis of the RA content of *Xenopus laevis* embryos during early gastrula estimated that they contain about 100-150 nM RA [71–77]. To perform physiologically-relevant manipulations of RA we increased that level by about 10%-25% using 10 nM and 20 nM treatments, respectively [27]. Embryos were treated from late blastula to early gastrula (st. 10.25) and collected for expression analysis by qPCR. This analysis revealed robust responses by the *dhrs3* homoeologs (p<0.0001) and weak (not significant) responses by the rest of the genes tested (Fig. 5). The self-regulation of the RA metabolic and signaling network to maintain or restore normal signaling levels is widely accepted, and the transcriptional responses of *aldh1a2, cyp26a1, dhrs3*, and *rdh10* to increased RA levels are the basis of this suggestion [18– 24,70,78]. In a recent study, we performed a detailed analysis of the RA responsiveness and requirement for the RA network genes expressed during early gastrula [27]. While some genes exhibited robust and concentration-dependent responses, others showed no significant changes in response to RA fluctuations. Also, the same gene was shown to exhibit different responsiveness at different developmental stages [27]. These changes in responsiveness could be explained in part by the different temporal expression patterns and the RA responsiveness of the homoeologs that were not addressed in previous studies.

**Figure 5.**
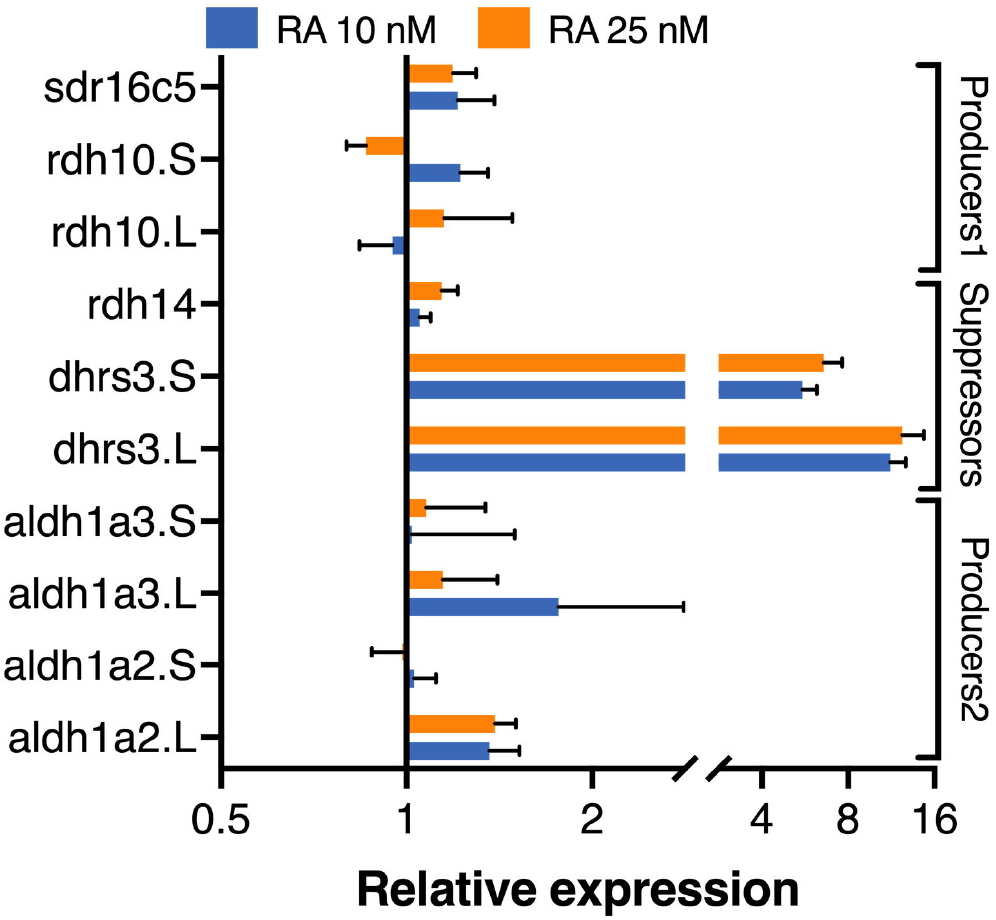
Responsiveness of homoeologs and singletons to RA manipulations. Embryos were treated from late blastula with 10 nM or 25 nM RA. Samples were collected at early gastrula (st. 10.25) and analyzed by qPCR for the RA responsiveness of individual homoeologs and singletons using the primers listed in Table.1. Expression changes were normalized to transcript levels in control embryos.

### 3.4. Homoeolog response to transient RA manipulation

The enhanced homoeolog gene loss observed within the RA metabolic network leading to the production of RAL raised the question as to the possible selective pressure driving this phenomenon. Maintenance of normal RA signaling levels is central for the prevention of the teratogenic effects of increased or decreased RA signaling levels [13,21,71,79–82]. Thus, homoeolog gene loss might be a “solution” to achieve tighter signaling regulation, i.e., robustness [27]. Several models can be suggested that could drive this gene loss to achieve higher RA signaling robustness. One possibility is that one of the homeologs in the pair has a less efficient or enhanced sensitivity, “noisier” regulation, and this gene is lost preferentially to reach tighter regulation. A “noisy” gene could mediate fast responses and might not necessarily be advantageous to lose. Alternatively, coordinated regulation of numerous genes performing the same enzymatic function might be more complicated to achieve, so having fewer genes would provide tighter regulation. To discriminate between these possibilities, we took advantage of an experimental protocol for transient RA manipulation and kinetic monitoring of the recovery process by qPCR (Fig. 6A) [27]. This assay allows us to monitor the robustness response as it takes place by analysing the expression changes of RA network components and downstream, RA-regulated genes. To perform physiologically relevant RA manipulations, based on our homoeolog responsiveness analysis (Fig. 5) and our previous studies [27], embryos were treated with 10 nM and 25 nM all-*trans* RA for 2 hours from late blastula to early gastrula (st. 10.25). The treatment was terminated by washing (T0) and samples were collected during the recovery period at 1.0, 1.5, 2.0, and 2.5 hours post-washing (T1, T1.5, T2, and T2.5, respectively) (Fig. 6A). RNA samples were prepared for comparative expression analysis to control samples. For enzymes that oxidize ROL we analyzed *rdh10*.L, *rdh10*.S, and *sdr16c5* (Producers 1), for enzymes that reduce RAL to ROL we chose *dhrs3*.L, *dhrs3*.S, and *rdh14* (Suppressors), and among the genes that produce RA (Producers 2), we studied both homoeologs of *aldh1a2* and *aldh1a3* (Fig. 1). For each homoeolog or singleton, the relative expression (fold change; •) was calculated at each time point relative to the expression in sibling control embryos at the same developmental stage, and the average • of four biological replicates was calculated. Analysis of *rdh10*.L, *rdh10*.S, and *sdr16c5* revealed very slight fluctuations of all three genes irrespective of whether 10 nM or 25 nM RA was used for the treatment (Fig. 6B,C). These weak responses are in agreement with the previous results of the RA responsiveness in which all three genes responded similarly (Fig. 5A). Also, these results suggest that none of the genes exhibits heightened responses in our experimental protocol which aims to mimic physiological RA fluctuations.

**Figure 6.**
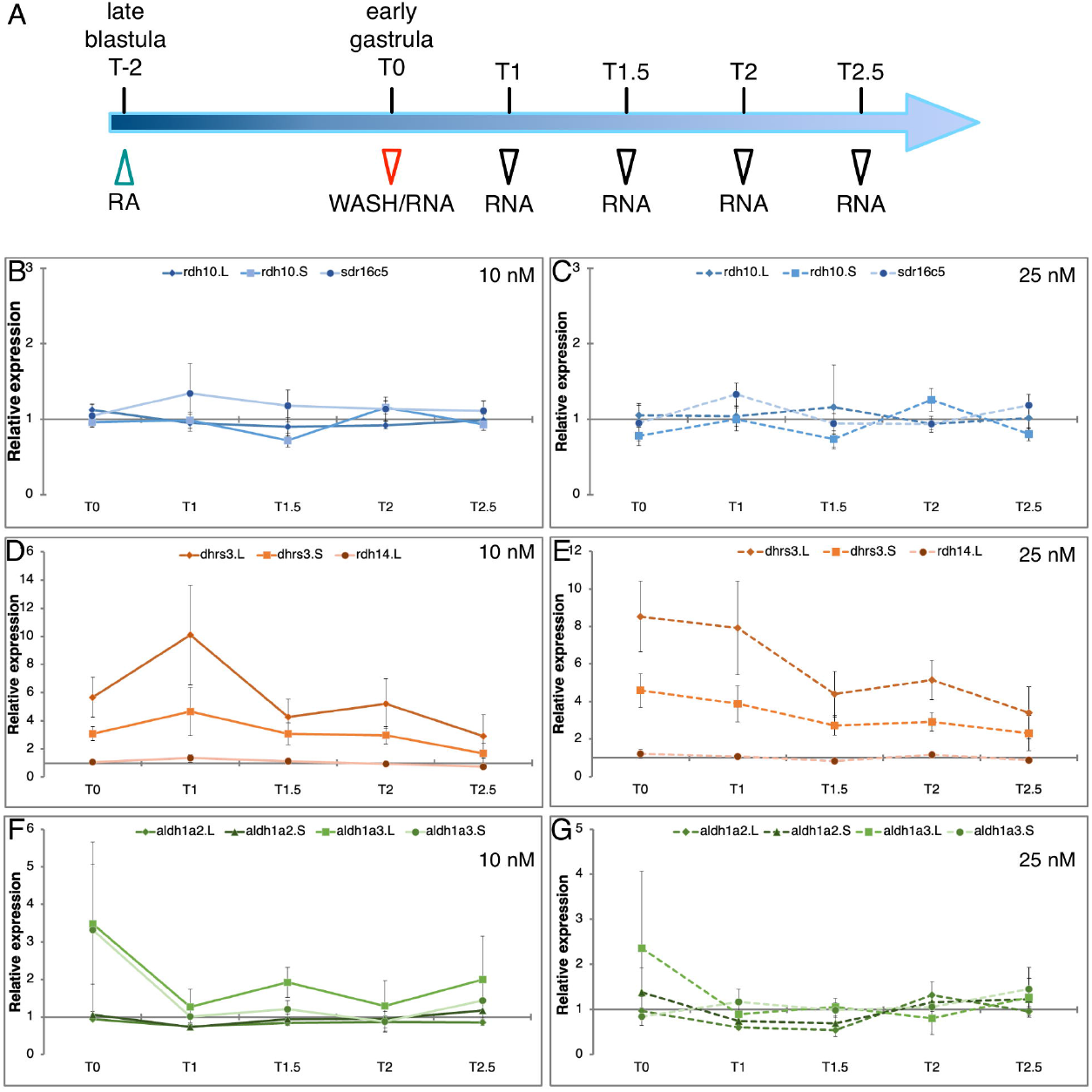
Recovery of RA metabolic gene expression following transient RA manipulation. (A) Embryos were subjected to a two-hour (T-2 - T0) RA treatment (10 nM or 25 nM) from late blastula (st. 9) to early gastrula (st. 10.25). At T0 the treatment was terminated (washed) and the embryos were further incubated. Samples were collected at different times (red and black arrowheads) for expression analysis. (B,C) Kinetic analysis of *rdh10*.L, *rdh10*.S, and *sdr16c5* expression changes. (D,E) qPCR analysis of the expression of *dhrs3*.L, *dhrs3*.S, and *rdh14*. (F,G) Analysis of the *aldh1a2*.L, *aldh1a2*.S, *aldh1a3*.L, and *aldh1a3*.S expression.

Analysis of the genes encoding enzymes preferentially involved in reducing RAL to ROL, *dhrs3*.L, *dhrs3*.S, and *rdh14*, revealed the hypothesized situation where one of the homoeologs exhibits an enhanced response to changes in RA levels and delayed restoration of normal expression levels. Expression of *dhrs3*.L shows the strongest up-regulation at T0 irrespective of the amount of RA employed of the three genes analyzed (Fig. 6D,E). By comparison, *dhrs3*.S shows a robust but weaker response at the end of the treatment, and *rdh14* only exhibits a weak response. Importantly, while *dhrs3*.S and *rdh14* reached almost normal expression levels (<2.3 fold) at the end of the recovery period (T2.5), *dhrs3*.L is still significantly up-regulated (>2.9 fold)(Fig. 6D,E). Analysis of the *aldh1a2* and *aldh1a3* homoeologs revealed that by the end of the transient treatment (T0), some of the homoeologs exhibit a clear up-regulation, but already one hour into the recovery period all genes are almost back to normal expression levels (Fig. 6F, G). These results show that the transient RA treatments can induce robust responses (*dhrs3*), but many of the genes studied exhibit mild expression changes and strong robustness responses, i.e., return to normal levels. A previous study that employed the same transient RA manipulation protocol but collected samples up to 5.5 hours after washing showed efficient recovery to normal expression levels of most RA network genes [27].

To make comparisons between samples, genes, or biological replicates easier, we calculated the fold change of each gene for all time points (•) and we summed up the values into a summary fold change score (••). This score should be low for tightly regulated genes and high for enhanced responders with slow recovery to normal values. We calculated this regulation tightness score for all genes studied (Table 4). The results show that for genes exhibiting moderate or limited gene responsiveness to RA and expression changes throughout the recovery period, *rdh10*.L, *rdh10*.S, *sdr16c5, rdh14, aldh1a2*.L, and *aldh1a2*.S, the average •• score ranged from 4.3 to 5.8 (Table 4). For each one of these genes, the difference in the amount of RA added had a very limited effect on the variation in their expression. For genes with higher fluctuation in their expression, *aldh1a3*.L, *aldh1a3*.S, *dhrs3*.L, and *dhrs3*.S, the •• score increased in correlation with the kinetic analysis result (Table 4; Fig. 6). The score for *dhrs3*.L reached ••>28 making it the least tightly regulated gene or the gene with the most extreme responses to RA fluctuations of those analyzed. Importantly, both singletons studied, *rdh14* and *sdr16c5* exhibited tight regulation with low responsiveness to RA fluctuations even though they have been shown to be involved *in vivo* in the metabolism of RA [83– 85].

**Table 4.** Regulation tightness score for the RA network homoeologs and singletons.

| Gene | $\Sigma \Delta$ score<sup>1</sup> | |
| --- | --- | --- |
|  | 10nM RA | 25nM RA |
| **Suppressors** |  |  |
| *dhrs3.L* | 28.1 | 29.4 |
| *dhrs3.S* | 15.5 | 16.4 |
| *rdh14.L* | 5.2 | 5.1 |
| **Producers 1** |  |  |
| *rdh10.L* | 4.9 | 5.2 |
| *rdh10.S* | 4.8 | 4.6 |
| *sdr16c5* | 5.8 | 5.3 |
| **Producers 2** |  |  |
| *aldh1a2.L* | 4.3 | 4.4 |
| *aldh1a2.S* | 4.9 | 5.2 |
| *aldh1a3.L* | 10.0 | 6.4 |
| *aldh1a3.S* | 7.8 | 5.5 |

<sup>1</sup>Regulation tightness score

### 3.5. RA responsiveness in homoeolog CRISPant embryos

The results of the individual homoeolog responses to transient RA manipulation identified gene pairs that represent all the possibilities initially suggested. The *rdh10* and *aldh1a2* genes have tightly and similarly RA-regulated homoeolog pairs. Similar but slightly enhanced responses were observed for the *aldh1a3* homoeologs, whereas the *dhrs3* homoeolog pair showed strong responses to RA changes and marked differences between the L and S genes. To begin to address the possible force driving the gene loss that gives rise to singletons, we took advantage of the CRISPR / Cas9 technology to create a partial, homoeolog-specific gene loss. We designed homoeolog-specific single guide RNAs (sgRNA) for the *dhrs3* and *rdh10* genes to knock down the expression of one homoeolog without affecting the second one. We also designed a sgRNA targeting the *sdr16c5* singleton. To determine the efficiency of the sgRNAs, DNA was extracted from CRISPant embryos and the genomic region containing the sgRNA targeted sequence was PCR-amplified and sequenced. Decomposition analysis of the sequence traces [50,51] provided a quantitative assessment of the genome editing efficiency. Analysis of the sequencing traces demonstrated the creation of indels around the sequence targeted by the sgRNA and the deterioration of the sequencing quality (Supplementary Fig. S1). According to the decomposition analysis, we observed a relatively robust effect of the sgRNAs inducing indels (Supplementary Fig. S1).

To study the effect of losing one of the homoeologs on the RA robustness response, embryos were injected at the one-cell stage with the appropriate sgRNA /Cas9 riboprotein complex to generate CRISPant embryos, which were then subjected to the transient RA manipulation protocol using low RA concentrations (10 nM)(Fig. 6A). RNA samples were collected at T0 (wash) and at 1.0, 1.5, 2.0, and 2.5 hours after treatment termination. To understand the effect of this gene knockdown on the RA signaling levels, we analyzed the expression of a panel of RA-regulated genes including *cyp26a1*.L, *hoxd1*.L /S, *hoxa1*.L, *hoxa1*.S, *hoxa2*.L / S, *hoxb4*.S, *hoxb1*.L, and *hoxb1*.S (Fig. 7 and Supplemental Fig. S2). Comparison at T0 of the expression levels of RA-responsive genes between RA treatment and *rdh10, dhrs3*, and *sdr16c5* CRISPants treated with RA supported the efficiency of the sgRNAs (Fig. 7A,B and Supplemental Fig. S2A,B). The RA treatment alone induced up-regulation of all RA targets ranging from 3.6 to 35 fold increase, while RA treatment of the *rdh10*.L, *rdh10*.S, and *sdr16c5* CRISPants resulted in a weaker RA-induced up-regulation irrespective of the gene being knocked down (Fig. 7A and Supplemental Fig. S2A). In agreement with the similar and limited responses to RA exposure (Fig. 6B,C), the three genes individually targeted, *rdh10*.L, *rdh10*.S, and *sdr16c5*, resulted in similar outcomes. This response of the RA target genes agrees with the suggestion that between similarly regulated RA network genes that encode enzymes performing the same metabolic reaction, their loss is equivalent in the early embryo. Maintenance of the singletons might reflect different spatial-temporal regulation to perform the same enzymatic reaction in different tissues in the embryo or the adult.

**Figure 7.**
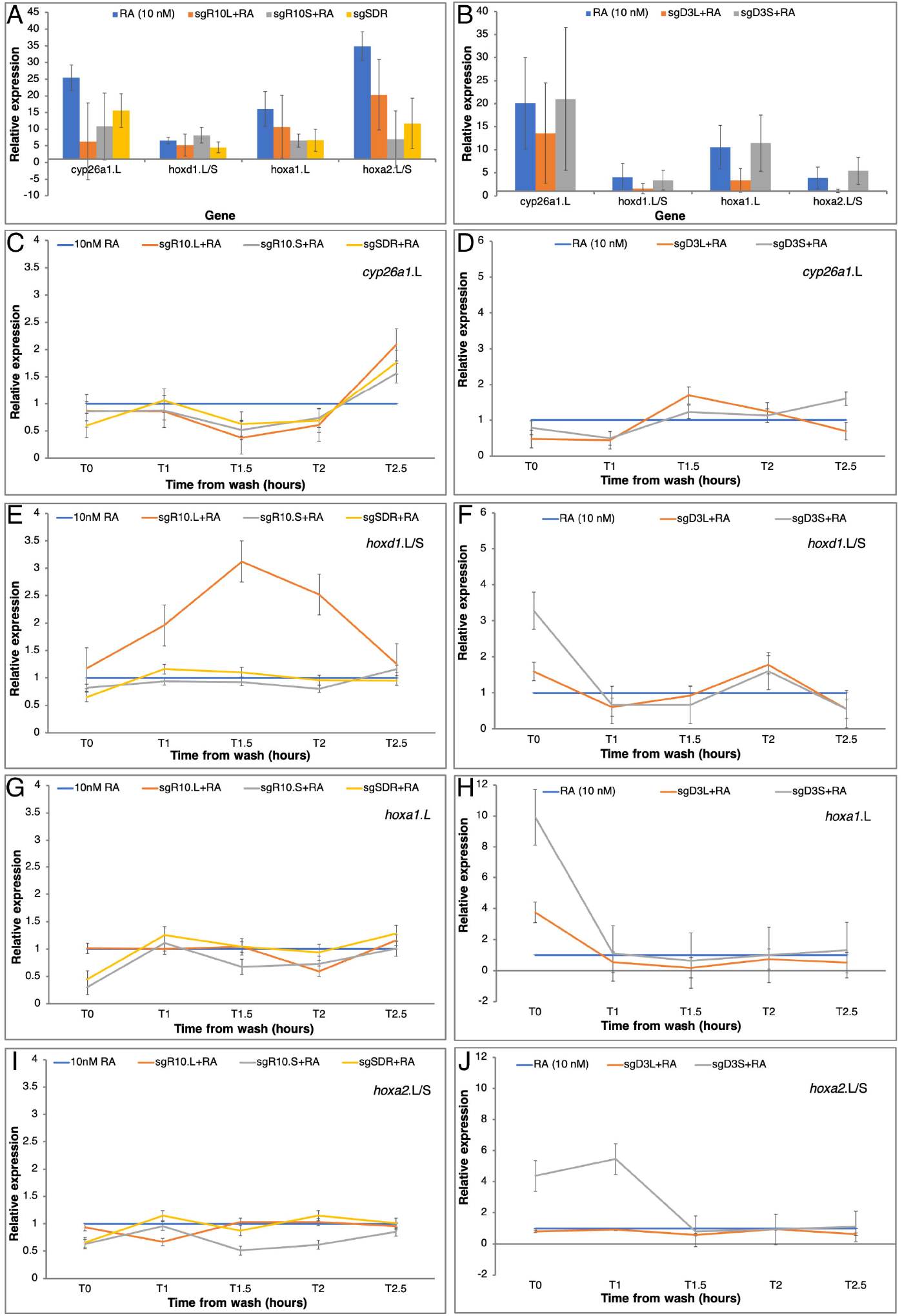
Gene expression changes in RA responsive genes as a result of homoeolog knockdown. RA network component gene specific knockdowns were induced by targeting genes with CRISPR /Cas9. The *rdh10*.L, *rdh10*.S, and *sdr16c5* (A,C,E,G,I), and *dhrs3*.L and *dhrs3*.S (B,D,F,H,J) genes were targeted with specific sgRNAs. CRISPant embryos were treated with RA (10 nM) and sibling embryos were treated with RA only as controls. (A,B) Gene expression change analysis at T0 normalized to control expression. (C-J) Kinetic analysis of gene expression changes in CRISPant embryos compared to RA-induced changes at each time point. Genes analyzed: (C,D) *cyp26a1*.L, (E,F) *hoxd1*.L / S, (G,H) *hoxa1*.L, (I,J) *hoxa2*.L /S.

Kinetic analysis of the RA robustness response in the *rdh10*.L, *rdh10*.S, and *sdr16c5* CRISPants was monitored by following the expression of the RA target genes during the recovery period (T0-T2.5; Fig. 6A). To better understand the contribution of the RA network components studied, the CRISPant samples treated with RA were compared to siblings treated with RA only. In most instances, knockdown of each of these three genes had a mild effect on the gene expression reducing the response to the transient RA treatment (Fig. 7C,E,G,I and Supplemental Fig. S2C,E,G,I). It is important to note that weaker responses in the CRISPants treated with RA support a tighter regulation of the signal as a result of the phenocopy of the gene loss. In a few instances we observed enhanced responses to the RA exposure mainly linked to the *rdh10*.L CRISPant (Fig. 7E and Supplemental Fig. S2C), the rest of the samples exhibited a more restricted response to RA exposure supporting a tighter regulation. To simplify the comparative analysis between CRISPants, we calculated the regulation tightness score (••) of the RA target genes for all time points compared to the response in RA-treated embryos (Fig. 8A). The scores for the RA-regulated genes in the three RA-treated CRISPants showed that all of them reduced the target gene expression changes. This result suggests that in the case of *rdh10*.L, *rdh10*.S, and *sdr16c5*, the gene activity reduction results in tighter regulation of the RA robustness response in agreement with a gene loss model for better regulation of the signal. Also in this analysis, we can observe the enhanced responses linked to the *rdh10*.L CRISPant (Fig. 8A).

**Figure 8.**
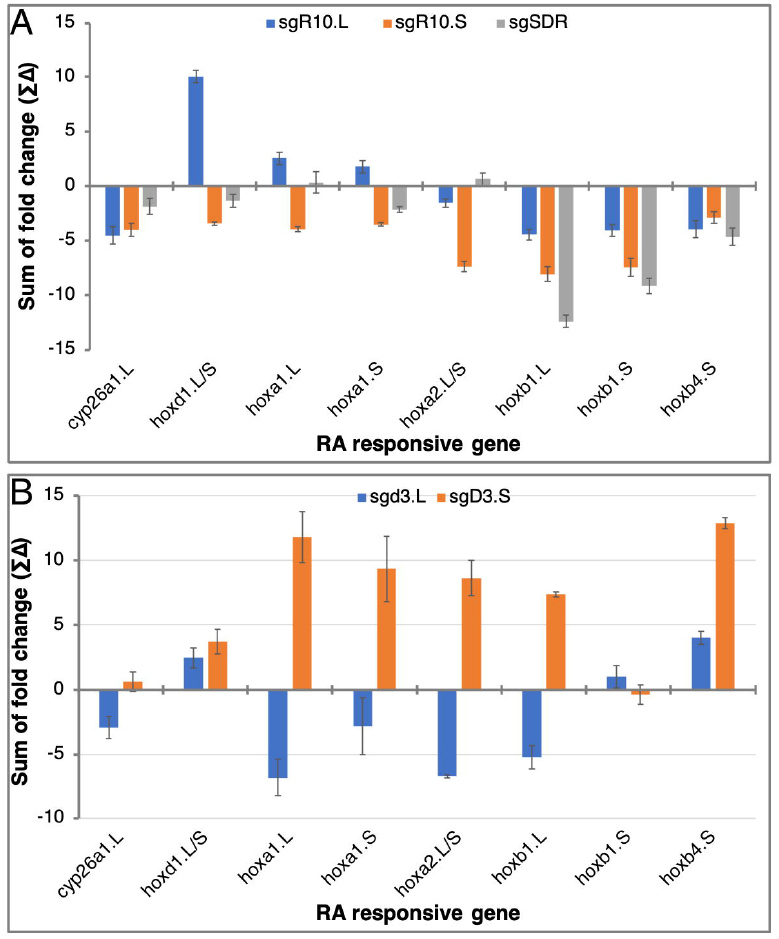
Regulation tightness score of the RA-responsive genes in CRISPant embryos. To calculate the regulation tightness score (••), the sum of the expression fold change was calculated for the RA-responsive genes: *cyp26a1*.L, *hoxd1*.L / S, *hoxa1*.L, *hoxa1*.S, *hoxa2*.L / S, *hoxb4*.S, *hoxb1*.L, and *hoxb1*.S. (A) Analysis in *rdh10*.L, *rdh10*.S, and *sdr16c5* RA-treated CRISPants. (B)Analysis in *dhrs3*.L and *dhrs3*.S RA-treated CRISPants.

The *dhrs3* homoeologs exhibit enhanced responses to the transient RA exposure with the *dhrs3*.L gene showing the strongest responses (Fig. 6D,E; Table 4). Since the DHRS3 enzyme preferentially reduces RAL back to ROL [25,59,86], the *dhrs3* CRISPants should exhibit enhanced RA signaling unless the RA network self regulation and robustness response compensates for this loss of activity [27]. Supporting the robustness scenario, the *dhrs3* CRISPants alone had almost no effect on the RA responsive genes with the exception of the two *hoxb1* homoeologs exhibiting a 1.5 to 7.5 increase in expression compared to control samples (not shown). Analysis at T0 of both *dhrs3* CRISPants treated with RA treatment showed that these responses were not enhanced as expected, and the partial knockdown of one of the *dhrs3* homoeologs resulted in reduced responses. In agreement with the loss of the loosely regulated homoeolog, these results show that the *dhrs3*.L CRISPants exhibit a stronger reduction in the RA response compared to knockdown of the *dhrs3*.S homoeolog (Fig. 7B and Supplemental Fig. S2B). Then, knockdown of the loosely regulated homoeolog achieves tighter regulation of the response.

While the T0 analysis supports the loss of the “noisier” gene to achieve tighter regulation of the response during the RA treatment, analysis of the full recovery kinetics compared to RA-only manipulated embryos provides information as to the effects of the homoeolog knockdown on the RA robustness response. Analysis of the same panel of RA-responsive genes showed that by about 1.5 hours into the recovery period (T1.5), the expression levels of the target genes tested in the *dhrs3*.L and *dhrs3*.S CRISPants was almost back to the same as the samples treated only with RA (Fig. 7D,F,H,J and Supplemental Fig. S2D,F,H,J). We could observe slight fluctuations in expression levels but in most instances, both CRISPants gave similar variations although the responses in the *dhrs3*.L tend to be lower than the RA-only samples, while the *dhrs3*.S CRISPants gave slightly enhanced responses. For multiple genes, at T0 we observed the up-regulation characteristic of the treatment before RA washing (Fig. 7F,H,J and Supplemental Fig. S2D,J). Also, at T0 and T1, CRISPants of the more tightly regulated homoeolog, *dhrs3*.S, exhibit larger fluctuations in expression of the RA responsive genes. Calculation of the regulation tightness score showed the opposed outcomes of the homoeolog-specific knockdown (Fig. 8B). The RA robustness response is enhanced by knockdown of the *dhrs3*.S homoeolog, while knockdown of *dhrs3*.L results in a reduced response. Analysis of the *dhrs3* homoeologs identified the *dhrs3*.L gene as the one exhibiting enhanced responses to RA treatment and less tight regulation (Fig. 6D,E). Then, knockdown of the homoeolog exhibiting tighter regulation exposes the system to the homoeolog with the gene with the apparent looser regulation. Interestingly, within 1.5 hours in the recovery, the system appears to stabilize irrespective of the homoeolog manipulated even though both *dhrs3* homoeologs take longer to reach normal expression levels. These observations suggest that the RA network robustness is capable of restoring normal signaling levels irrespective of the homoeolog knocked down.

## 4. Conclusions

The RA metabolic and signaling network is strongly dependent on the nutritional status and is influenced by the environment. Fluctuations in the RA signaling levels during embryogenesis can result in severe teratogenic outcomes. The preferential gene loss of the RA network components involved in the metabolism leading to RAL production suggests a selective pressure to achieve tighter regulation of the robustness response. Eliciting a RA robustness response by transient manipulation together with knockdown of specific genes support the suggestion that gene loss is linked to more efficient regulation of the network. Tighter network regulation might involve loss of homoeologs similarly regulated, or homoeologs with enhanced responses. While the allotetraploid condition of *X. laevis* is convenient to explore these genomic changes and their regulatory outcomes, in diploid organisms, besides gene duplications and deletions, mutation, addition and deletion of regulatory elements might take place to achieve the same outcome.

## Supplementary Materials

Supplementary Figure S1. Genomic sequence deterioration in the RA network CRISPants.

Supplemental Figure S2: RA responsiveness in RA network homoeolog-specific CRISPants.

## Acknowledgments

We wish to thank Martin Blum and Tim Ott for introducing us to the manipulation of the Xenopus genome using CRISPR /Cas9. We also wish to thank Sally Moody and Graciela Pillemer for critically reading the manuscript.

## Author Contributions

Conceptualization, A.F., L.B.-K. and T.A.; Methodology, T.A.; Validation, T.A., L.B.-K. and A.S.; Formal Analysis, T.A., L.B.-K. and A.S.; Investigation, T.A., L.B.-K. and A.S.; Resources, X.X.; Data Curation, T.A., L.B.-K. and A.S.; Writing – Original Draft Preparation, A.F.; Writing – Review & Editing, T.A., L.B.-K. and A.S.; Supervision, A.F.; Project Administration, A.F.; Funding Acquisition, A.F.”,

## Institutional Review Board Statement

Experiments were performed after approval and under the supervision of the Institutional Animal Care and Use Committee (IACUC) of the Hebrew University (Ethics approval no. MD-17-15281-3).

## Informed Consent Statement

Not applicable

## Data Availability Statement

Data is contained within the article or supplementary material

