## Supplemental Fig. S1 for "Enhanced loss of retinoic acid network genes in *Xenopus laevis* achieves a tighter signal regulation"

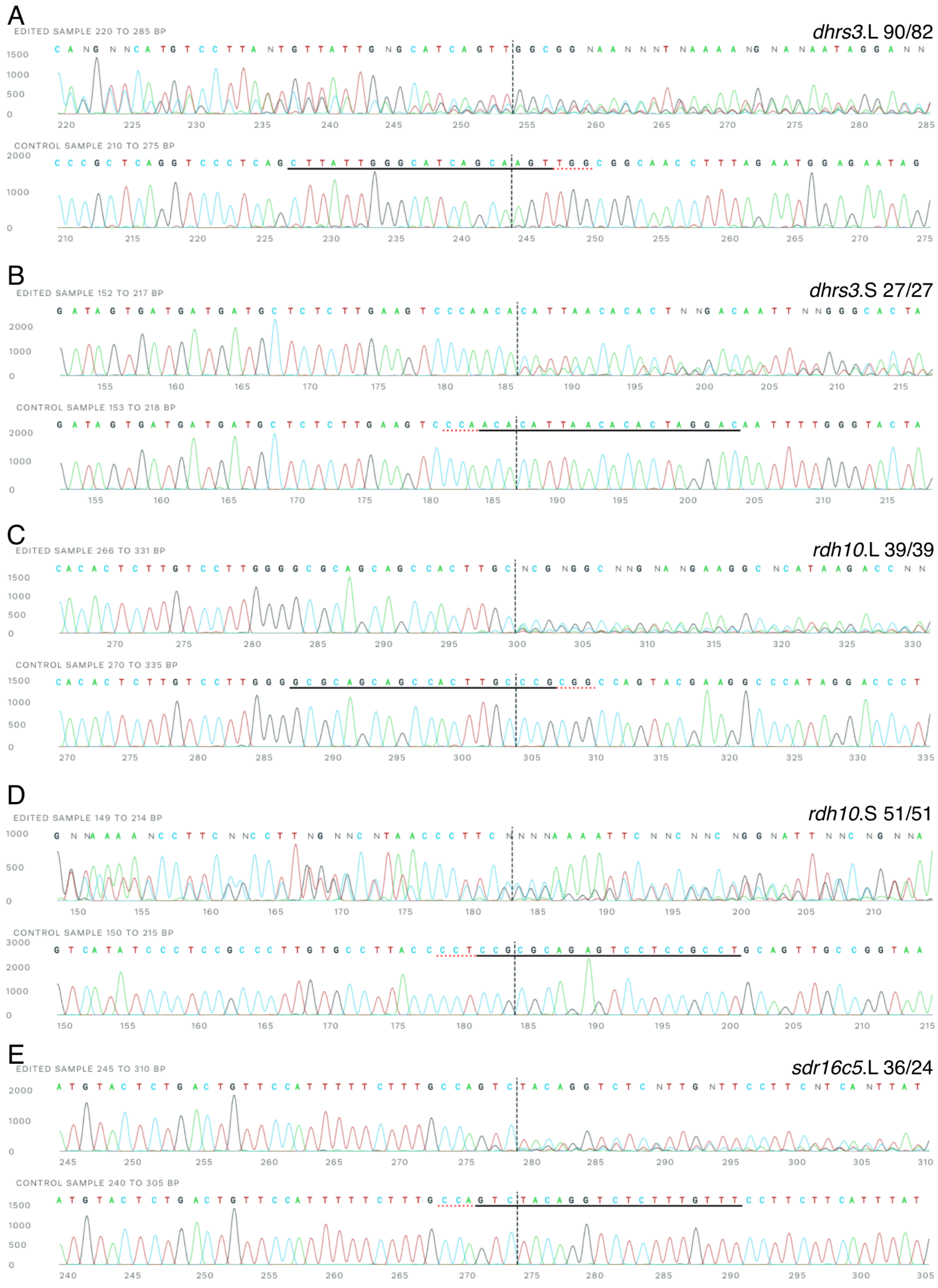


**Supplementary Figure S1. Genomic sequence deterioration in the RA network CRISPants.** CRISPant embryos were processed for DNA extraction and PCR amplification of the genomic region targeted by the sgRNA. Control embryos were processed in parallel for sequence comparison. For the *dhrs3*.L (A), *dhrs3*.S (B), *rdh10*.L (C), *rdh10*.S (D), *sdr16c5* (E) CRISPants, the corresponding control sequence is shown underneath. The horizontal line marks the position of the sgRNA in the sequence including the PAM site. The numbers after the gene name represent the indel and frameshift efficiencies from the decomposition analysis.
