## Supplemental Fig. S2 for "Enhanced loss of retinoic acid network genes in *Xenopus laevis* achieves a tighter signal regulation"

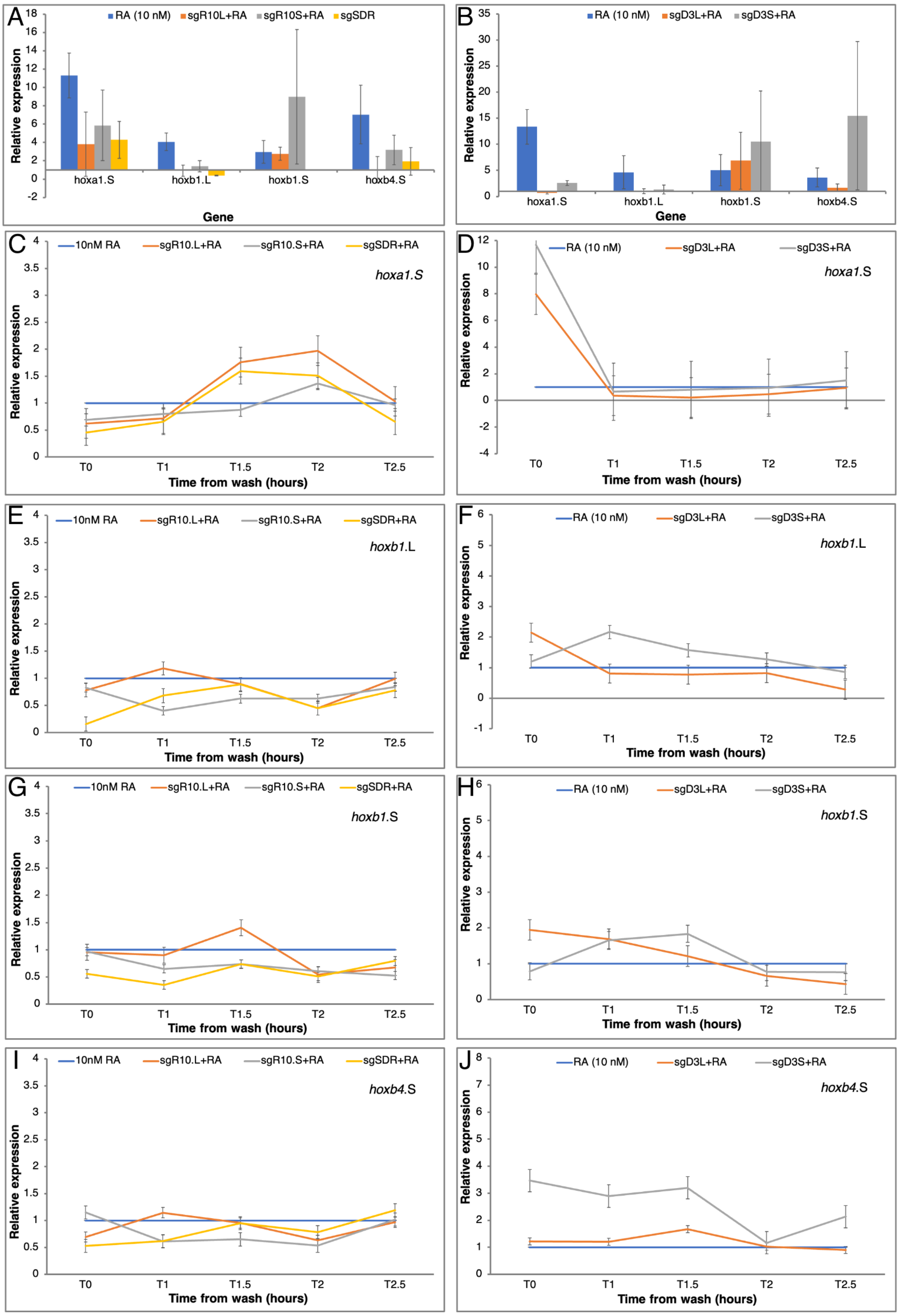


**Supplemental Figure S2. RA responsiveness in RA network homoeolog-specific CRISPants.** RA network component gene specific knockdowns were induced by targeting genes with CRISPR/Cas9. The *rdh10*.L, *rdh10*.S, and *sdr16c5* (A,C,E,G,I), and *dhrs3*.L and *dhrs3*.S (B,D,F,H,J) genes were targeted with specific sgRNAs. CRISPant embryos were treated with RA (10 nM) and sibling embryos were treated with RA only as controls. (A,B) Gene expression change analysis at T0 normalized to control expression. (C-J) Kinetic analysis of gene expression changes in CRISPant embryos compared to RA-induced changes at each time point. Genes analyzed: (C,D) *hoxa1*.S, (E,F) *hoxb1*.L, (G,H) *hoxb1*.S, (I,J) *hoxb4*.S.
